# Exercise-conditioned tear fluid suppresses myopia progression

**DOI:** 10.64898/2026.02.05.704049

**Authors:** Huilin Yao, Maodi Liang, Qintong Fei, Jing Cao, Tiantian Liang, Xiangtian Zhou, Sen Zhang, Qinghua Cui

**Affiliations:** School of Sports Medicine, Wuhan Sports University, No. 461 Luoyu Rd. Wuchang District, Wuhan 430079, Hubei Province, China; State Key Laboratory of Eye Health, Oujiang Laboratory (Zhejiang Lab for Regenerative Medicine, Vision and Brain Health), Eye Hospital, Wenzhou Medical University, Wenzhou, China; Department of Biomedical Informatics, State Key Laboratory of Vascular Homeostasis and Remodeling, School of Basic Medical Sciences, Peking University, 38 Xueyuan Rd, Beijing, 100191, China

**Keywords:** myopia, exercise, tear fluid, reverse transcriptomics

## Abstract

Myopia represents a major global public health challenge with rapidly rising prevalence. It is thus important to explore novel therapeutics for the treatment of myopia. Here we predicted tear fluid after (TA) but not before (TB) moderate-intensity aerobic exercise might protect against myopia using reverse transcriptomic analysis. To experimentally validate this hypothesis, TA or TB was administered by periocular injection in a guinea pig model of form-deprivation myopia. As a result, TA treatment significantly attenuated myopic refractive shifts and suppressed vitreous chamber depth and axial length elongation, whereas TB showed no protective effects. This study proposes a novel therapeutic avenue for myopia intervention and also suggests a previously unrecognized tear fluid–mediated mechanism linking exercise to myopia.

## 1 Introduction

Myopia is a major global public health challenge, with a rapidly increasing prevalence worldwide, particularly in East Asia^[1-2]^. Epidemiological studies estimate nearly half the global population may be affected by 2050, with many progressing to high myopia and facing vision-threatening complications such as retinal detachment, myopic maculopathy, and glaucoma^[3-5]^. Myopia is structurally characterized by excessive axial elongation, driven by coordinated changes in multiple ocular tissues, including the retina, choroid, and sclera^[6-8]^. Although optical and pharmacological interventions—such as orthokeratology lenses and low-dose atropine—can slow myopia progression, their use is limited by patient compliance, side effects, or uncertain long-term efficacy^[9-11]^. Therefore, identifying novel therapeutics remains an important research goal.

Lifestyle factors play a critical role in the development and progression of myopia. Among these, exercise is one of the most consistently reported protective factors against myopia onset and progression in children^[12-14]^, and this protective effect is largely attributed to enhanced retinal dopamine release^[15-16]^. Beyond this local mechanism, exercise also induces systemic physiological changes in cardiovascular, metabolic, immune, and nervous systems, and growing evidence suggests that these systemic effects may also influence ocular physiology^[17-19]^.

Tear fluid is an easily accessible biological fluid that reflects both local ocular surface conditions and broader physiological states. Previous tear research has shown that exercise can increase tear secretion, improve tear film stability, and reduce levels of inflammatory markers in tear fluid, particularly in individuals with dry eye disease^[20-22]^. Proteomic analyses have further revealed that tears contain a wide array of bioactive molecules, including growth factors, cytokines, enzymes, and neuromodulators, which can influence cell behavior and tissue homeostasis^[23-27]^. Despite these insights, existing research has largely concentrated on changes in tear volume or composition, however, the biological effects of tear fluid and their potential regulation by systemic physiological states such as exercise remain underexplored.

Reverse transcriptomics represents a powerful tool to explore potential indications for a given ‘drug’ agent by comparing the gene signatures induced by the drug with those of diseases^[28-29]^. Specifically, if a drug induces gene expression changes that are inversely correlated with those observed in a disease state, the drug may have therapeutic potential on this disease. Critically, this strategy bypasses the need to dissect individual components of complex bioactive body fluids (e.g., tear fluid), making it ideal for evaluating the collective functional effects of heterogeneous bioactive molecules.

In this study, we first identified the gene signatures induced by tear fluid from healthy adults after (TA) and before (TB) moderate-intensity aerobic exercise by analyzing the RNA-sequencing data of their perturbed A549 cells. Next, using reverse transcriptomics, we systematically analyzed the relationships between the gene signatures of TA/TB and those of various diseases. The results showed that the TA-induced gene signature is significantly anti-correlated with those of myopia, while TB show no significance. This prediction suggest that TA could have protective effects on myopia, while TB could not have similar effects. To test this hypothesis, we conducted a proof-of-concept validation using a form-deprivation myopia (FDM) animal model, demonstrating that TA indeed significantly delayed myopia progression *in vivo*. These findings propose a novel therapeutic avenue for myopia intervention and also suggests a previously unrecognized tear fluid–mediated mechanism linking exercise to myopia.

## 2 Material and Methods

### 2.1 Study Participants and Ethics Approval

This study recruited five healthy adult female participants (aged 22-30 years, BMI 18-25 kg/m^2^) with no history of systemic diseases or ocular conditions (such as dry eye disease, ocular trauma) who had not used any medications affecting tear secretion or inflammatory responses in the past month. Participants underwent comprehensive pre-clinical screening. Basic personal information was recorded (Supplementary File S1), and all participants completed the Physical Activity Readiness Questionnaire (PAR-Q)^[30]^. Comprehensive pre-clinical screening was also performed, including basic ocular examinations such as visual acuity and intraocular pressure measurements, as well as blood pressure assessment and routine blood tests. These measures were implemented to ensure each participant could safely complete a standardized moderate-intensity aerobic exercise protocol. The study protocol was approved by the Ethics Committee of Wuhan Sports University (Approval No. 2025089; Approval Date: April 10, 2025), and written informed consent was obtained from all participants.

### 2.2 Exercise Intervention Protocol

Since none of the participants had regular exercise habits, we used the Tanaka formula to estimate their maximum heart rate: maximum heart rate = 208 − (0.7 × age)^[31]^. Based on this estimated value, the target exercise intensity was set at 60%–80% of the individual’s maximum heart rate, and the target heart rate range for each participant was calculated to be approximately 116–138 beats per minute^[21]^.

To control for confounding factors, all experimental sessions were scheduled to avoid the participants’ menstrual periods. 24 hours prior to the exercise intervention, participants were required to strictly adhere to a standardized dietary protocol: abstain from alcohol, coffee, strong tea, and other caffeinated beverages; and avoid high-sugar, high-fat, spicy and other stimulant-rich foods. This was to minimize the influence of dietary variations on metabolic state. Additionally, participants were instructed not to wear contact lenses for 24 hours before the experiment.

The specific exercise protocol was as follows: Participants first performed a 5-minute progressive warm-up on a treadmill, followed by 30 minutes of continuous running at their individual target heart rate^[21]^. Heart rate was monitored in real-time throughout the entire session using a Polar H10 heart rate sensor.

### 2.3 Tear Collection and Sample Processing

Tear samples were collected before and immediately after exercise. Tears were collected using sterile Schirmer test strips (5 mm wide), which were placed at the outer one-third of the lower conjunctival sac until full saturated. All procedures were performed by uniformly trained personnel to minimize variability^[32]^. Each strip was immediately transferred to a pre-cooled 2 ml centrifuge tube containing 500 μl of normal saline. The strips were then eluted on an orbital shaker at 4 °C for 2 h, followed by centrifugation at 4 °C and 4000 rpm for 5 min. The supernatant was collected and filtered through a 0.22 μm membrane. The total protein concentration was determined using BCA protein assay kit (Servicebio, Catalog No. G2026), and all samples were adjusted to a uniform concentration of 600 μg/ml with normal saline. Samples were aliquoted and stored at -80 °C until use.

### 2.4 Cell Culture and Treatment

In this study, A549 cells were used as the cell model for identifying the tear-induced gene signatures, as they exhibit robust and reproducible gene expression responses to diverse external perturbations ^[33-34]^. A549 cell lines were cultured in high-glucose Dulbecco’s Modified Eagle Medium (DMEM) supplemented with 10% fetal bovine serum (FBS) and 1% penicillin– streptomycin, and maintained at 37 °C in a humidified atmosphere containing 5% CO_2_. Logarithmically growing cells were seeded into 6-well plates. After overnight attachment, the medium was replaced with fresh medium supplemented with tear samples (TB or TA, 600 μg/mL), followed by incubation for 48 h. Three independent biological replicates were performed for each group.

### 2.5 RNA isolation, cDNA library preparation and sequencing

Total RNA was extracted with Trizol (Invitrogen) and assessed with Agilent 2100 BioAnalyzer (Agilent Technologies, Santa Clara, CA, USA) and Qubit Fluorometer (Invitrogen). Total RNA samples that meet the following requirements were used in subsequent experiments: RNA integrity number (RIN) > 7.0 and a 28S:18S ratio > 1.8.RNA-seq libraries were generated and sequenced by CapitalBio Technology (Beijing, China). The triplicate samples of all assays were constructed an independent library, and do the following sequencing and analysis. The NEB Next Ultra RNA Library Prep Kit for Illumina (NEB) was used to construct the libraries for sequencing. NEB Next Poly(A) mRNA Magnetic Isolation Module (NEB) kit was used to enrich the poly(A) tailed mRNA molecules from 1 μg total RNA. The mRNA was fragmented into ∼200 base pair pieces.

The first-strand cDNA was synthesized from the mRNA fragments reverse transcriptase and random hexamer primers, and then the second-strand cDNA was synthesized using DNA polymerase I and RNaseH. The end of the cDNA fragment was subjected to an end repair process that included the addition of a single “A” base, followed by ligation of the adapters. Products were purified and enriched by polymerase chain reaction (PCR) to amplify the library DNA. The final libraries were quantified using KAPA Library Quantification kit (KAPA Biosystems, South Africa) and an Agilent 2100 Bioanalyzer. After quantitative reverse transcription-polymerase chain reaction (RT-qPCR) validation, libraries were subjected to paired-end sequencing with pair end 150-base pair reading length on an Illumina NovaSeq sequencer (Illumina).

### 2.6 RNA-sequencing and data analysis

The genome of human genome version of hg38 was used as reference. The sequencing quality were assessed with FastQC (v0.11.5) and then low quality data were filtered using NGSQC (v2.3.3)^[35]^ .The clean reads were then aligned to the reference genome using HISAT2 (v2.1.0) with default parameters^[36]^. The processed reads from each sample were aligned using HISAT2 against the reference genome. The gene expression analyses were performed with StringTie (v1.3.3b)^[37]^. DESeq (v1.28.0) was used to analyze the differentially expressed genes (DEGs) between samples^[38]^. Thousands of independent statistical hypothesis testing was conducted on DEGs, separately. Then a p-value was obtained, which was corrected by FDR method. And Corrected P-value (q-value) was calculated by correcting using BH method. Statistical significance was determined based on adjusted p-values (false discovery rate, FDR), with q ≤ 0.05 considered significant. Parameters for classifying significantly DEGs are ≥2-fold differences (|log2FC|≥1, FC: the fold change of expressions) in the transcript abundance and p ≤ 0.05. The annotation of the DEGs were perfomed based on the information obtained from the database of ENSEMBL, NCBI, Uniprot, GO, and KEGG. The RNA-sequencing data are available at GEO (GSE315463).

### 2.7 Prediction of potential protective effects of the tear after exercise

Based on the gene expression profiles induced by control agent, tear before exercise (TB), and tear after exercise (TA), We first calculated fold change of TA vs control, TB vs. control, and TA vs. TB. To investigate the potential biological TA and TB, the genes exhibiting a fold change (FC) ≥ 1.50 or ≤ 0.67 were designated as the transcriptomic gene signatures. Next, using an in-house *in silico* platform and reverse transcriptomics-based screening program, we predicted the potential diseases the tear could treat. The hypothesis is that the diseases whose gene signatures are inversely correlated with the tear’s gene signatures could be treated by tear.

### 2.8 Experimental Animals and Housing Conditions

This study utilized three-week-old tricolor pigmented guinea pigs (Cavia porcellus, English shorthair stock, n=35). The animals were housed under a 12/12-hour light/dark cycle at a controlled room temperature of 20-26℃. Cage floor illuminance was maintained at approximately 300 lux. All guinea pigs had ad libitum access to standard chow and fresh vegetables, which were provided twice daily. The experimental protocol was approved by the Animal Care and Ethics Committee of Wenzhou Medical University (Wenzhou, China) and conformed to the ARVO Statement for the Use of Animals in Ophthalmic and Vision Research.

### 2.9 Ocular Biometry

Refraction measurements were performed in a dark room. Refractive status was assessed without cycloplegia using a custom-built infrared photorefractor, measuring the vertical pupil meridian as previously described^[39]^. Three readings were taken per eye and averaged for analysis. Axial length (AL) was measured using an A-scan ultrasonograph (11 MHz, AVISO Echograph Class I-Type Bat; Quantel Medical, France). Prior to measurement, topical anesthesia was induced with a drop of 0.5% proparacaine hydrochloride (Alcon, Belgium). Ten consecutive traces were captured per eye, and the mean value was recorded as the final AL.

### 2.10 Statistical Analysis

All datasets were confirmed to follow a normal distribution with equal variance. Data are presented as mean ± standard deviation. Interocular differences (treated eye minus untreated fellow eye) for refractive error and AL were compared across groups using repeated-measures ANOVA, with treatment group as the between-subjects factor and time as the within-subjects factor. Post-hoc analyses were conducted with Bonferroni correction. Statistical significance was set at p < 0.05. All analyses were performed using GraphPad software (version 10.6.0; GraphPad Software, LLC, USA).

## 3 Results

### 3.1 Diseases predicted to be treated by tear after exercise

The overall workflow of this study is shown in Figure 1. Tear samples were collected from healthy adults before (TB) and after (TA) moderate-intensity aerobic exercise and used to treat A549 cells for 48 h (600 μg/mL). Transcriptomic profiling was then performed to define gene expression signatures induced by TB and TA. Reverse transcriptomic analysis was performed to evaluate disease relevance by testing for inverse correlations between tear-induced gene signatures and disease-associated gene signatures using Fisher’s exact test. As a result, for the gene signature of TA-control, 26 diseases (Supplementary File S2) were filtered out using the cutoff of odds ratio (OR) <1 and p-value<0.05. Using the same procedure, 40 diseases and 24 diseases were filtered out for TA vs. TB, and TB vs. control, respectively (Supplementary File S2). More interestingly, for diseases are significant both in TA vs. control and TA vs. TB (Figure 2). Among which, both predictions suggest that TA could have therapeutic effects on myopia.

**Figure 1.**
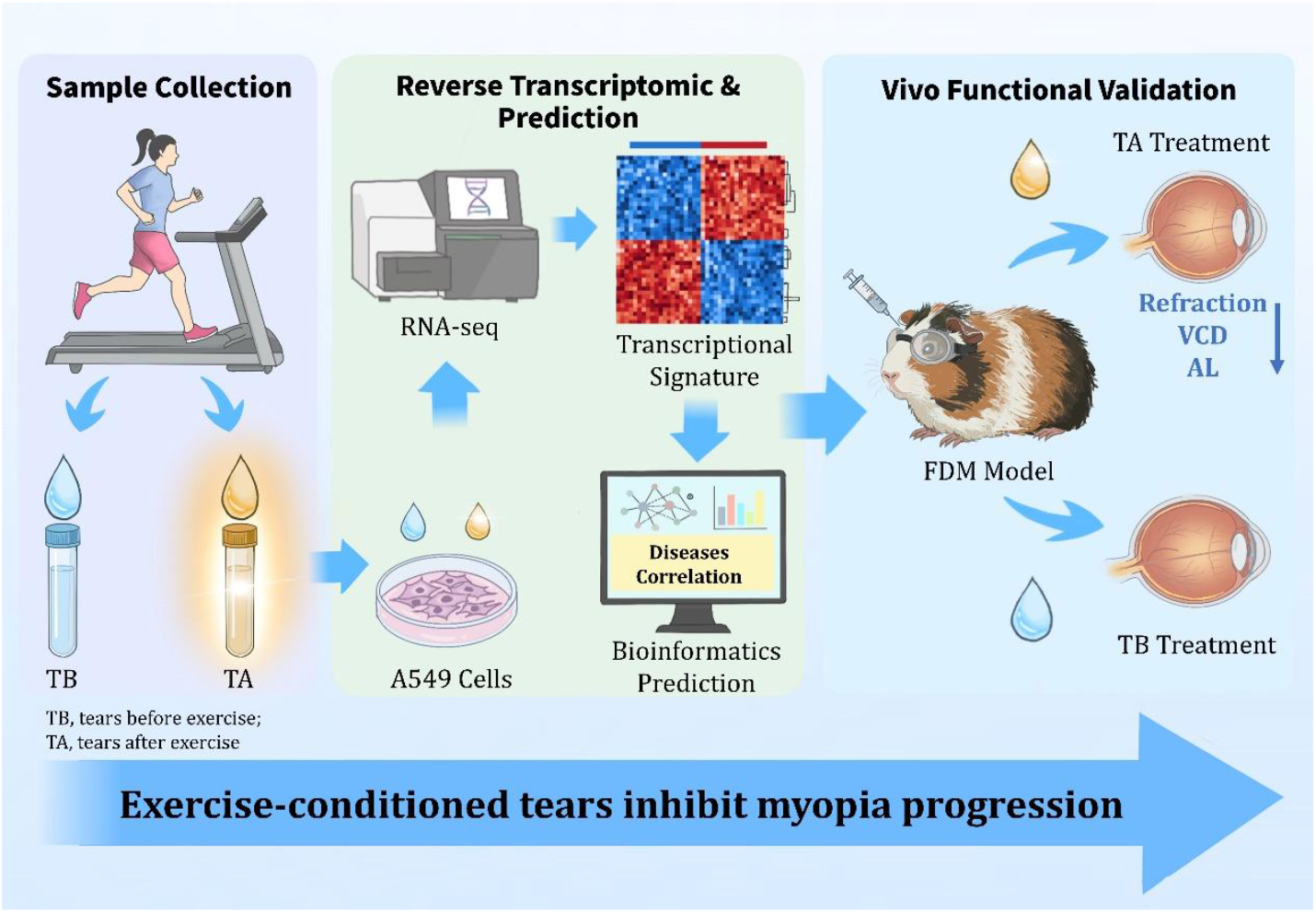
Overview of the study design and experimental workflow. TB, tears before exercise; TA, tears after exercise.

**Figure 2.**
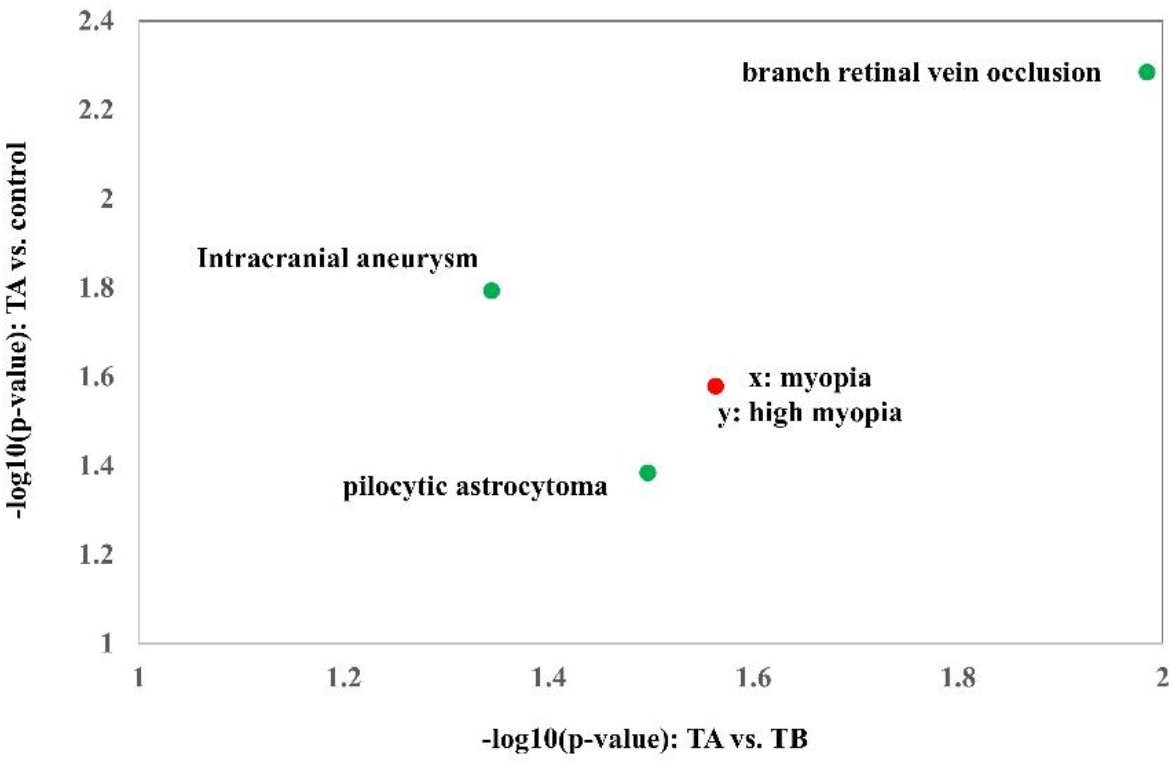
Reverse transcriptomic prediction of diseases potentially modulated by tear after exercise.

### 3.2 Tear after exercise (TA) suppresses myopia progression in vivo

To assess the potential therapeutic effect of TA on myopia, we examined their effects following myopia induction via the form-deprivation (FD) procedure (Supplementary Figure S1). Periocular injection of TA and TB elicited differential effects on myopia progression. Repeated measures ANOVA revealed a significant main effect of treatment on the myopic shift in refraction (F_(2,32)_=6.775, p < 0.01), vitreous chamber depth (VCD) elongation (F_(2,32)_=7.194, p < 0.01), and axial length (AL) elongation (F_(2,32)_=11.57, p < 0.001). Eyes treated with TA exhibited a significantly smaller myopic shift compared to vehicle-treated FDM controls. This inhibitory effect was significant at the 1-week timepoint (TA: -2.40 ± 1.05 D vs. Vehicle: -4.31 ± 1.26 D; p < 0.001) and maintained at 2 weeks post-injection (TA: -4.27 ± 1.26 D vs. Vehicle: -5.79 ± 1.49 D; p < 0.01, Figure 3A). Refractive changes in the TB group, however, closely mirrored the vehicle control at both time points, demonstrating no therapeutic benefit. Consistent with the refractive data, TA treatment significantly attenuated the abnormal axial elongation characteristic of myopia development. The suppression of VCD elongation was significant at both 1 and 2 weeks (both p < 0.01, Figure 3B). Similarly, the increase in AL was markedly reduced in the TA group compared with the vehicle control at both time points (both p < 0.001, Figure 3C). In contrast, TB treatment failed to alter the progression of axial elongation, with changes in VCD and AL indistinguishable from the vehicle control. Collectively, these results indicate that TA effectively inhibit myopia progression by limiting both refractive shift and axial elongation.

**Figure 3.**
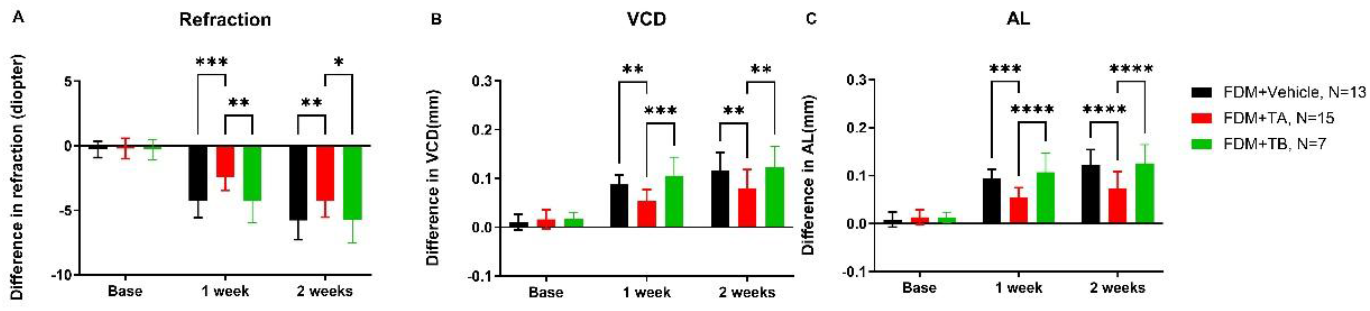
Tear after exercise suppresses FDM progression in vivo. Form-deprivation myopia (FDM) was induced in guinea pigs, followed by periocular injection of vehicle, tear fluid collected before exercise (TB), or tear fluid collected after exercise (TA). (A) Interocular differences in refraction (treated eye minus fellow eye) at baseline, 1 week, and 2 weeks after treatment. TA-treated eyes exhibited a significantly smaller myopic shift compared with vehicle-treated FDM controls, whereas TB treatment showed no significant effect. (B) Interocular differences in vitreous chamber depth (VCD). TA significantly attenuated VCD elongation at both 1 and 2 weeks, while TB did not differ from vehicle controls. (C) Interocular differences in axial length (AL). TA treatment markedly suppressed abnormal axial elongation associated with myopia development, whereas TB failed to alter AL progression.Data are presented as mean ± SD. Statistical significance was determined using repeated-measures ANOVA followed by Bonferroni post hoc tests. *p < 0.05, **p < 0.01, ***p < 0.001, ****p < 0.0001. Group sizes were as follows: FDM + Vehicle, n = 13; FDM + TA, n = 15; FDM + TB, n = 7.

## Discussion

As a major global public health concern, myopia leads to sight-threatening complications like retinal detachment, myopic maculopathy, and glaucoma. While optical and pharmacological interventions are available, significant challenges remain in achieving effective and sustainable myopia control. In this study, we demonstrated that tear fluid collected after exercise (TA) exhibits distinct biological activity compared with that collected before exercise (TB), and that this activity is functionally relevant to the regulation of myopia. By integrating reverse transcriptomic prediction with in vivo validation, we found that exercise-conditioned tear fluid induces a unique gene expression signature and significantly suppresses the progression of FDM in guinea pigs. These results provide a new perspective for understanding how exercise-induced systemic physiological states influence ocular axial growth through tear fluid-mediated mechanisms.

Tear fluid is a complex biological liquid containing proteins, lipids, electrolytes, metabolites, and various signaling molecules, which collectively participate in regulating ocular surface and epithelial cell behavior^[24-26]^. Growing evidence indicates that tears are not merely lubricating secretions—their contained growth factors, cytokines, and neuromodulators can regulate cell proliferation, inflammatory responses, and tissue remodeling^[23-27]^. Previous research on exercise and ocular physiology has primarily focused on tear secretion volume, tear film stability, and ocular surface inflammatory status. Studies have shown that aerobic exercise can increase tear secretion and prolong tear film break-up time, an effect particularly evident in individuals with dry eye disease^[20-22]^. Additionally, acute exercise can reduce the levels of pro-inflammatory cytokines and oxidative stress markers in tear fluid^[21]^. Although these studies suggest that exercise can alter tear composition, they did not further explore whether such compositional changes are accompanied by alterations in biological function. Our study further revealed that exercise can induce qualitative changes in the bioactivity of tear fluid, enabling TA to exert biological effects that modulate disease-relevant physiological processes. Thus, the shift from compositional changes to enhanced biological activity in exercise-conditioned tears underscores their potential not only as indicators of physiological status but also as a source of bioactive factors for therapeutic applications in myopia and other ocular disorders.

A notable strength of this study lies in the application of reverse transcriptomic analysis to infer the protective effects of tear fluid. This strategy has been widely used to identify interventions that reverse disease-associated gene expression signatures^[28-29]^. Its advantage is that it does not require prior knowledge of specific active molecules, enabling hypothesis-free functional evaluation of complex biological perturbations. By treating tear fluid as a functional biological perturbation, we screened for disease types whose transcriptional signatures were inversely correlated with those induced by TA. Notably, myopia was consistently predicted as a potential target in both the TA vs. control and TA vs. TB comparisons, whereas TB showed no such predictive association. This indicates that exercise induces a substantive transformation in tear fluid function, rather than merely amplifying its baseline activity. Subsequent in vivo experiments strongly supported the biological relevance of these transcriptomic predictions. The *in vivo* experiments provided direct evidence for the suppressive effect of TA on myopia progression. In a FD-induced myopia model in guinea pigs, periocular administration of TA significantly alleviated myopic refractive changes and inhibited increases in vitreous chamber depth and axial length, while the TB-treated group showed no similar protective effect. Given that excessive axial elongation is a primary structural basis for the development of myopia^[6-8]^, these results suggest that TA may act by interfering with core regulatory mechanisms of axial growth, rather than producing only transient optical effects.

The development of myopia involves coordinated signaling among the retina, retinal pigment epithelium, choroid, and sclera, ultimately leading to remodeling of the scleral extracellular matrix and axial elongation^[40-42]^. Although the specific molecular components responsible for the effects of TA remain to be identified, it is plausible that exercise-modulated tear constituents contribute to ocular growth regulation by influencing inflammatory responses, oxidative stress, or growth factor-related signaling pathways. These biological processes are well established as key regulators of scleral remodeling and axial elongation during myopia development^[43-44]^. The ability of TA to simultaneously suppress refractive error and axial elongation further supports its genuine disease-modifying role.

Epidemiological studies consistently indicate that exercise reduces the risk of myopia onset and progression in children^[12-14]^. It is widely accepted that enhanced light exposure and retinal dopamine release are primary protective mechanisms^[15-16]^. However, the role of exercise-related systemic factors remains incompletely understood. This study proposes that tear fluid may serve as a potential mediator linking physical activity to axial growth regulation, offering a complementary mechanism to existing light-dependent pathways. Unlike environmental light exposure, tear-mediated mechanisms may exert local and paracrine effects at the ocular surface and could potentially influence deeper ocular tissues through neural or biochemical signaling pathways. This perspective expands current understanding of the relationship between exercise and myopia protection and emphasizes the active role of body fluids in physiological regulation.

Several limitations of this study should be noted. First, the human sample included only young adult females, it remains unclear whether the TA of males and aged or younger also have similar effects. Second, it is also unclear whether the findings keep consistent for other modes of exercises. Moreover, reverse transcriptomic analysis primarily identified functional associations and cannot pinpoint the specific tear components responsible for the observed effects. Currently, the identical bioactive molecules and its mechanisms on preventing myopia remain unknown. Future work should integrate proteomics, metabolomics, and functional fractionation techniques to identify key bioactive molecules and their mechanisms. Furthermore, although periocular injection provided proof-of-concept in this study, clinically feasible delivery strategies remain to be explored.

## Conclusions

In summary, this study demonstrates that exercise induces functional changes in tear fluid, which can be effectively identified through reverse transcriptomic analysis and validated by in vivo experiments. Tear after exercise (TA) significantly slowed myopia progression, with concomitant reductions in vitreous chamber and axial elongation. These findings establish a novel link between exercise, tear biology, and the regulation of axial growth, suggesting that tear fluid as a previously unrecognized mediator of exercise-induced myopia protection. Importantly, this mechanism may provide new avenues for understanding and potentially intervening in myopia.

## Acknowledgements

This study was supported by the Noncommunicable Chronic Diseases-National Science and Technology Major Project (2024ZD0531201) and Natural Science Foundation of China [82427801, 32301239, 62025102] and the Scientific and Technological Research Project of Xinjiang Production and Construction Corps (2023AB057) and the Natural Science Foundation of Hubei Province (2025AFD622).

## Completing interests

The authors declare no competing interests.

## Author Contributions

QC proposed the project. XZ and SZ designed the animal experiments. HY, ML, and QF performed experiments of RNA-sequencing and reverse transcriptomics. JC and TL took part in data analysis. SZ performed the animal experiments. HY, SZ, and QC wrote the draft manuscript. XZ, SZ, and QC supervised the study.

## Supplementary Information

**Supplementary File S1**. Baseline characteristics of participants.

**Supplementary File S2**. Disease signatures significantly inversely correlated with tear-induced gene expression profiles.

---

**Supplementary Figure S1.**
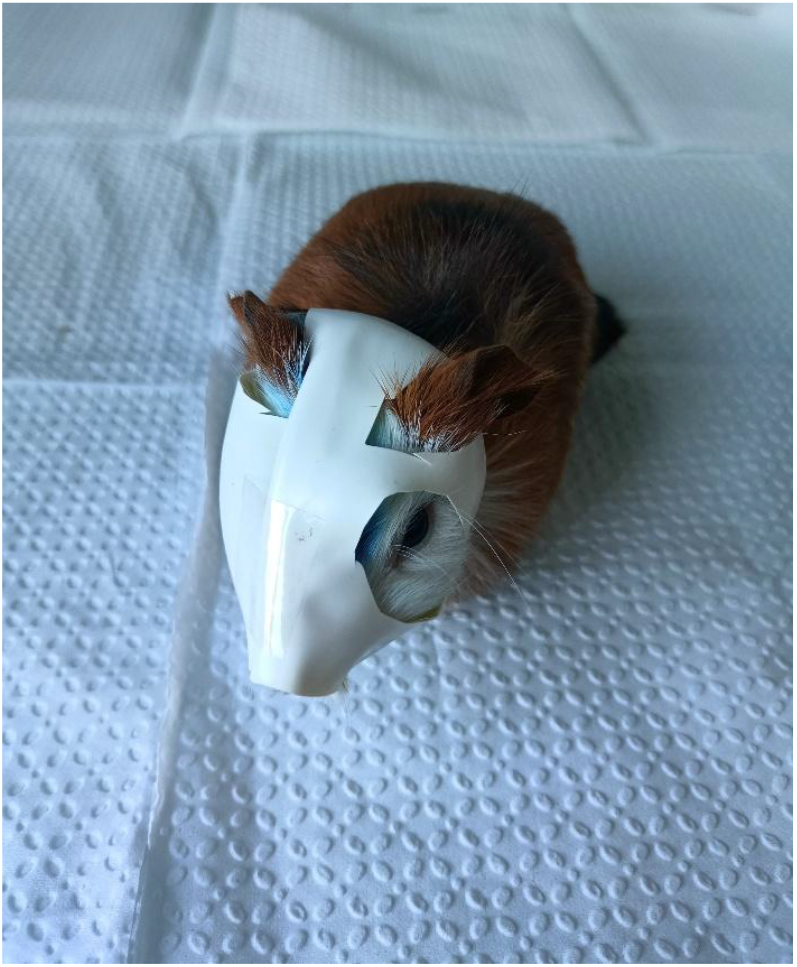
Establishment of the form-deprivation myopia (FDM) model.

